# The prevalence of protein misfolding as a mechanism for hereditary deafness

**DOI:** 10.64898/2026.03.09.710547

**Authors:** Rose A. Gogal, Genevieve M. Cox, Diana L. Kolbe, Amanda M. Odell, Chloe E. Ovel, Katherine I. McCormick, Brian Hong, Hela Azaiez, Thomas L. Casavant, Richard J.H. Smith, Terry A. Braun, Michael J. Schnieders

## Abstract

Hearing loss is the most common sensory deficit impacting ∼5% of the world’s population. The Deafness Variation Database (DVD) is a public resource of deafness variants, containing over 380,000 missense variants across 224 genes, with 303,577 classified as a variant of uncertain significance (VUS). To address the challenge of evaluating each deafness associated VUS, we evaluate a family of probabilistic frameworks to quantify the strength of computational evidence based on ACMG/AMP recommendations. First, CADD and REVEL are compared using Bayesian models parameterized using either a ClinVar 2019 dataset or labeled DVD variants. The REVEL model built using the DVD dataset demonstrates the best accuracy, sensitivity, and specificity. Incorporation of (in)tolerance to missense variation based on sorting each gene into three bins (tolerant, average, intolerant) shows that intolerant DVD genes are consistent with a higher prior probability of being pathogenic (25.7%) than average (10.7%) or tolerant (8.7%) genes. Finally, the impact of protein folding stability was incorporated using a 2D likelihood, which surpassed the simpler models while also offering a biophysical rationale for the disease mechanism. The protein folding-informed Bayesian model results in 28,866 prioritized VUSs reaching a posterior probability of pathogenicity above 98% with a false positive rate of only 0.14%. Overall, 54,752 missense variants are predicted to cause protein folding destabilization of greater than 1.0 kcal/mol, while 18,706 of the 28,886 prioritized VUS (65%) surpass this threshold. From these VUSs, we identify twelve probands where the patient’s genetic diagnosis is upgraded to likely pathogenic/pathogenic. We highlight two variants that cause clear structural disruption, demonstrating the impact of biophysical characterization on variant evaluation.

**Author Summary:** We investigate the impacts of single amino acid changes on protein structure and folding in the context of hearing loss. Hearing loss is the most common impairment of the main senses affecting nearly 5% of the world’s population. About 45% of people with hearing loss receive a diagnosis after targeted genetic testing. Here, we integrate biophysical data that quantifies the effect of a change to protein sequence on protein folding in combination with genetic data to improve our ability to identify protein amino acid changes that are likely to impact hearing. Our work leads to 12 patients receiving an upgraded diagnosis with their variant disrupting protein stability. Although the method is applied to hearing loss, it can be used for interpreting protein sequence changes in other disease contexts.

## Introduction

Hearing loss is the most common sensory deficit. It affects ∼5% of the world’s population, impacting people of all ages and exacting significant personal and societal costs.(1) Amongst newborns, 1 to 3 of every 1000 babies have hearing loss, with about 60% of cases due to genetic factors, and by the age of 80, 50% of octogenarians will required some form of auditory amplification for meaningful communication(1, 2). Although a Mendelian inheritance pattern is common for genetic hearing loss, extreme genetic and phenotypic heterogeneity drive an underlying complexity that makes the genetic diagnosis of hearing loss challenging(3). Typically, establishing a genetic diagnosis necessitates multidisciplinary expertise, with input from otolaryngologists, human geneticists, genetic counselors, bioinformaticians, biomedical engineers, and auditory scientists(3).

A genetic diagnosis guides the clinical care of persons with hearing loss. For that reason, the OtoSCOPE^®^ sequencing panel was developed at the University of Iowa in 2012 to provide comprehensive genetic testing for hearing loss. Version 9 targets 224 genes associated with non-syndromic and syndromic hearing loss(4) and has a diagnostic rate across all patients of 45%(5). Variants identified by OtoSCOPE^®^ testing are classified following guidelines established by the American College of Medical Genetics and Genomics (ACMG) and Association for Molecular Pathology (AMP) into one of five categories: benign, likely benign, likely pathogenic, pathogenic, and variant of uncertain significance (VUS)(6) by combining varying levels of evidence (sequencing data, *in vivo* and/or *in vitro* experiments, family history, *etc.*) into a comprehensive framework.

The Deafness Variation Database(7) (DVD) is a publicly available database of 10,586,514 variants for OtoSCOPE^®^ genes that have been collected from dbSNP(8), gnomAD(9), 1000 Genomes Project(10), ClinVar(11) and HGMD(12). Of these variants, 668,489 are within exons or impact canonical splice sites, including 381,924 missense variants. These missense variants are classified as benign (0.75%, 2,895), likely benign (17.2%, 65,585), likely pathogenic (0.83%, 3,205), and pathogenic (1.74%, 6,662), with 79.5% (303,577 variants) being a VUS. Due to the intractable challenge of generating experimental evidence to assess the impact of VUSs, here we explore computational methods to identify missense VUSs that exhibit features consistent with pathogenicity.

Two popular *in silico* methods for large dataset variant evaluation are the Combined Annotation Dependent Depletion (CADD) and Rare Exome Variant Ensemble Learner (REVEL) scores. CADD evaluates the deleteriousness of all genetic variants based on a set of 63 genomic features that include conservation and regulatory annotations(13, 14). REVEL is an ensemble method, assessing the deleteriousness of missense variants by combining outputs of 13 individual tools, several of which use features like those in CADD(15). While CADD is a single model applicable to all variant types, REVEL’s ensemble approach is optimized for pathogenicity prediction in missense variants. Importantly, neither CADD nor REVEL model the structural or energetic consequences of variants on protein structure. However, both provide a numerical score to indicate the relative likelihood that a variant is deleterious *without* assigning an ACMG variant classification code.

Work by Pejaver et al.(16) has derived score thresholds for genetic effect predictors using Bayesian statistics. These score thresholds are consistent with varying levels of evidence following ACMG PP3/BP4 criteria where the lowest level of evidence (supporting) reaches a posterior probability of pathogenicity of 10% and the highest level of evidence (very strong), reaches a posterior probability of pathogenicity of 98%(16). Using the ClinVar 2019 dataset, Pejaver et al. found that no tool was able to provide a very strong evidence designation.

The structure of a protein dictates its function and phenotype. When evaluating the impact of a missense variant that causes a downstream amino acid change, it is critical to access the impact on protein folding stability(17). Folding free energy difference (*ΔΔG*_*Fold*_) quantifies the change in stability of a folded protein state upon mutation. Positive folding free energy differences indicate a destabilizing mutation thereby impacting function. With the introduction of AlphaFold3(18) (an improved protein structure prediction tool), the Bayesian framework introduced by Pejaver et al., and more than 380,000 missense variants in the DVD, we can combine genetic tools with protein folding information to help classify variants of uncertain significance. Previous work investigating the impact of disease-causing missense variants on protein folding found that ∼60% to ∼83% cause destabilization(19–21). We hypothesize that combining genetic tools with protein folding information for the specific phenotype of hearing loss will improve the predictive power of computational tools.

Here, we re-evaluate the thresholds proposed by Pejaver et al. in the context of hearing loss to improve the predictive power of CADD and REVEL. We use scores describing the tolerance of a gene to mutation from gnomAD to refine our classification methods by organizing genes into tolerance-based groups with specific priors(22). We combine the Bayesian framework for genetic effect predictors (*e.g.,* CADD and REVEL) with folding free energy differences to define a protein folding-informed classification approach with improved predictive power for prioritizing missense VUSs in the DVD as deleterious(16, 23). The clinical impact of this work is highlighted by the identification of 12 variants identified by OtoSCOPE^®^ that have very strong evidence for being pathogenic based on protein folding-informed modeling, and describe two variants in detail where the combination of genetic tools and protein folding stability demonstrate how the prediction of variant effect can be missed with genetic tools alone.

## Results

### Monomer Structures

The DVD provides structures of 258 proteins across 224 genes, decreasing the average MolProbity score from 1.85 to 1.05 after optimization with the AMOEBA force field (referred to as OtoProteinV3). The MolProbity scores for OtoProteinV3 are summarized in Figure 1A. Most structures have a MolProbity score between 0.5 and 1.0, as compared to the majority AlpaFold3 structures that fall between 2.0 and 2.5. The van der Waals clash score was reduced from 5.29 clashes per 1000 atoms to 0.15 clashes per 1000 atoms for OtoProteinV3. A comparison of OtoProteinV2 and OtoProteinV3 alpha tectorin structure (TECTA) is shown in Figure1B together with MolProbity statistics, reflecting the reported differences in accuracy for AlphaFold2 and AlphaFold3(18). Fig1C shows the significant (4-fold) increase in DVD missense variants investigated from OtoProteinV2 (2023) and OtoProteinV3 (2026).

**Figure 1.**
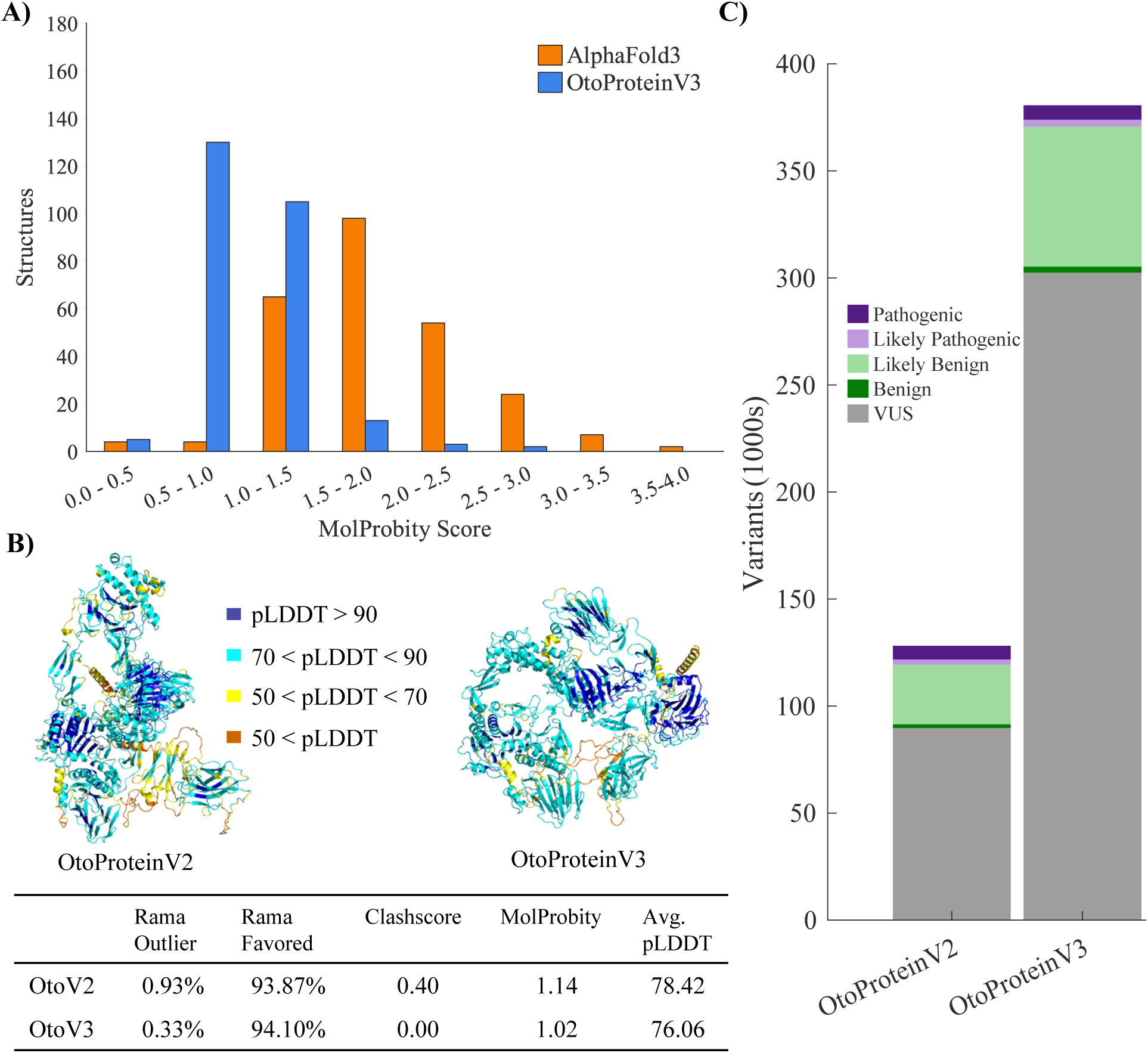
Summary of the structures and variants. Fig1A summarizes the MolProbity scores for raw AlphaFold3 structures (orange) and OtoProteinV3 optimized structures (blue). Fig1B shows an example of the OtoProteinV2 (2023) and OtoProteinV3 (2026) TECTA full-length protein structure colored by AlphaFold pLDDT confidence score. The MolProbity statistics for each structure are shown in the corresponding table. Fig1C shows the increase in the number of variants from the 2023 study to the 2026 study.

### Tolerance in Deafness-Associated Genes

Tolerance-based bins of DVD genes as compared to genome-wide tolerance were roughly comparable, with median missense o/e scores being 0.89 and 0.88, respectively. However, the o/e score distribution for the DVD skewed slightly more towards tolerant genes (skewness of DVD - 0.36 vs of genome -0.31).

We highlight three genes across the range of o/e scores in the DVD: *ACTG1*, *COCH*, and *USH1C*. *ACTG1* (Fig2A) is the second most intolerant gene in the DVD (the most tolerant being *WFS1*) with an o/e of 0.43. Gamma actin is highly conserved across species, and mutations in actin proteins are often lethal during development (selected against) due to their role in the structural framework of cells(24). Consequently, the structure of gamma actin is well-defined, which leads to more regions of high AlphaFold3 confidence (Fig2A). *COCH* encodes for the protein cochlin and has an o/e of 0.86. While cochlin is essential for hearing function, mutations are not as strongly selected against during development as those in gamma actin, and it has both well-folded regions and disordered regions, which contribute to the average tolerance (Fig 2B). *USH1C* (Fig2C) encodes for harmonin and pathogenic variants can be associated with autosomal recessive non-syndromic hearing loss or autosomal recessive Usher Syndrome Type I(25). *USH1C* has an o/e of 1.0 and its recessive inheritance pattern is consistent with its above average o/e. The structure for *USH1C* has the lowest level of predicted AlphaFold3 confidence.

**Figure 2.**
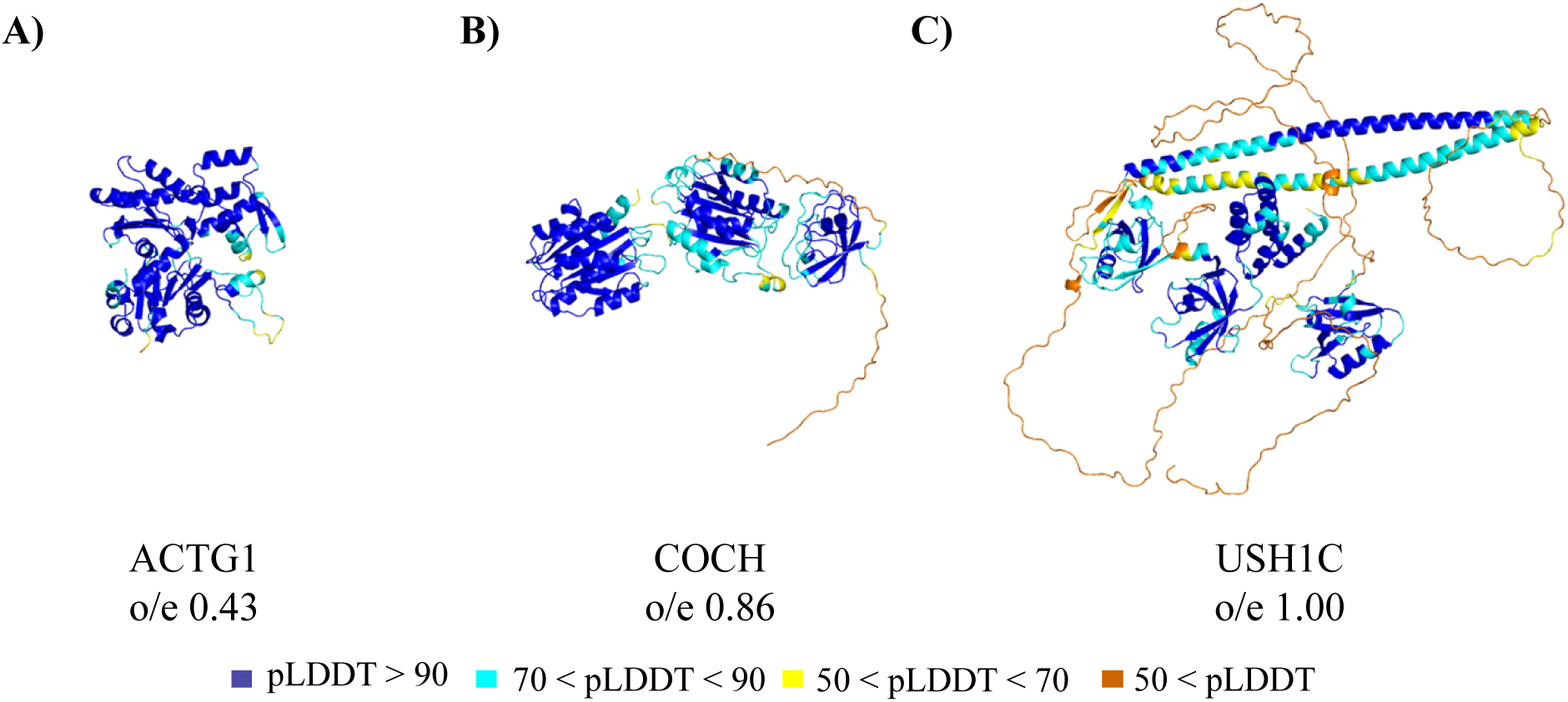
Three proteins across the o/e spectrum in the DVD colored by their AlphaFold3 confidence. Fig2A shows ACTG1, a gene within the intolerant bin with an o/e of 0.43. Fig2B shows COCH, an average tolerance gene that encodes for the protein Cochlin. Fig2C USH1C encodes for harmonin whose pathogenic variants are associated with Usherin Syndrome type I.

### Tolerance-Based Priors vs a DVD Prior

The ClinVar 2019 dataset resulted in a prior probability of pathogenicity of 4.1% (prior odds of 0.046) for their dataset, which is low compared to previous recommendations and genotype/phenotype prior calculations for similar Bayesian based studies(26, 27). Previous studies on the Chinese Deafness Consortium defined priors ranging from 9.31-28.5%(28). For DVD missense variants with a classification (∼80,000), the rate of pathogenicity is 12.6% (prior odds of 0.144), which is consistent with these other estimates.

We calculated tolerance-based priors by grouping genes using their o/e scores into three bins based on the distribution of o/e in the DVD, except for assigning X-linked/MT genes (where o/e scores are not available) to a separate bin. For each bin, Table 1 shows the o/e score range, the number of genes and variants (total and labeled), and the tolerance-based prior probability of pathogenicity/benignity. The bin for average tolerance contains most of the DVD genes with its center near the average missense o/e (± 0.1) for both the entire genome and subset of genes in the DVD. The tolerant and intolerant bins contain genes on the extreme ends of the DVD o/e range. The prior in the intolerant bin is much higher than that for the DVD prior, while that for the tolerant bin is lower. This supports the hypothesis that missense variants in genes highly intolerant to mutation are more likely to be pathogenic than are missense variants in genes tolerant to mutation.

**Table 1.**
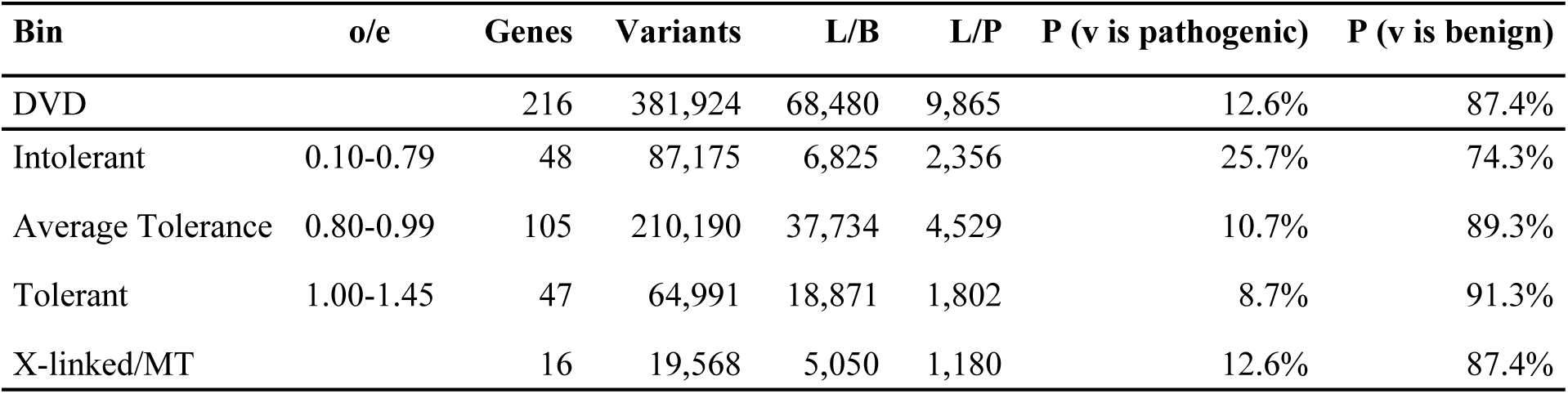
Description of the tolerance-based bins, genes, variants, and prior probabilities. The tolerance-based bins used for the DVD and resulting prior probabilities of pathogenicity and benignity as described in the text. o/e is the range of tolerance scores for each bin. Gene and variants are the number of genes and total variants that fall in each bin. L/B and L/P are the number of labeled variants in each bin. P (v is pathogenic) and P (v is benign) are the prior probabilities that a variant is pathogenic or benign in each bin. The X-linked/MT bin is all genes without an o/e score, which is assigned the overall DVD prior.

| Bin | o/e | Genes | Variants | L/B | L/P | P (v is pathogenic) | P (v is benign) |
| --- | --- | --- | --- | --- | --- | --- | --- |
| DVD |  | 216 | 381,924 | 68,480 | 9,865 | 12.6% | 87.4% |
| Intolerant | 0.10-0.79 | 48 | 87,175 | 6,825 | 2,356 | 25.7% | 74.3% |
| Average Tolerance | 0.80-0.99 | 105 | 210,190 | 37,734 | 4,529 | 10.7% | 89.3% |
| Tolerant | 1.00-1.45 | 47 | 64,991 | 18,871 | 1,802 | 8.7% | 91.3% |
| X-linked/MT |  | 16 | 19,568 | 5,050 | 1,180 | 12.6% | 87.4% |

### Genetic Tool Analysis

We calculated the posterior associated with REVEL or CADD and determined the score threshold for each evidence level (supporting, moderate, strong, or very strong). The threshold curve for the genetic tools and the posterior along the range of scores using a DVD likelihood and prior for both CADD and REVEL are shown in Figure 3. Fig3A and Fig3B show the REVEL posterior curve for benign and pathogenic variants, respectively. Fig3C and Fig3D show the CADD posterior curve for benign and pathogenic variants, respectively. We derived DVD and tolerance-based thresholds as displayed in Table 2.

**Figure 3.**
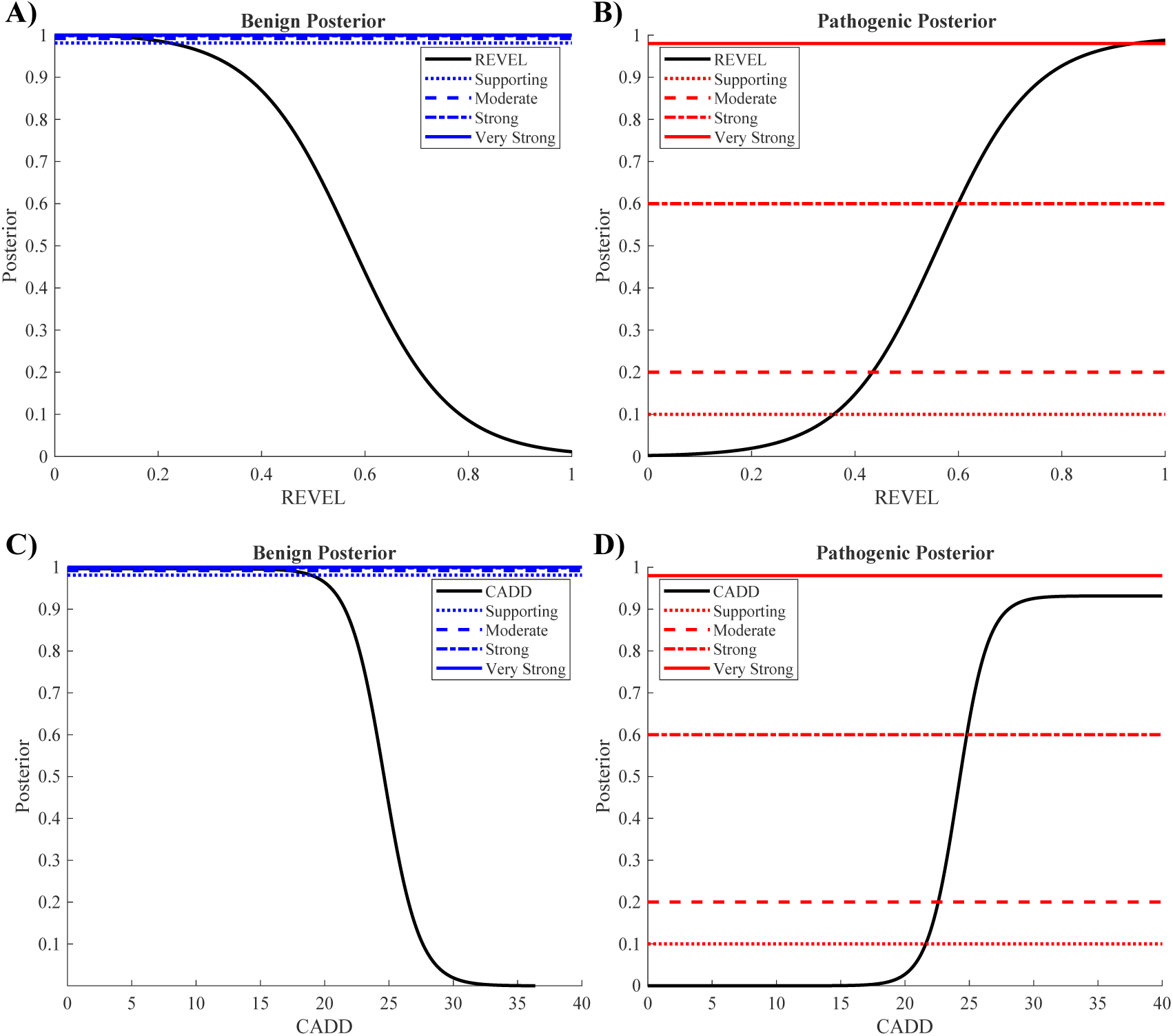
Posterior probability versus score plots using DVD derived likelihood and prior probability of pathogenicity. For each level of evidence (supporting, moderate, strong, very strong), dotted lines indicate the corresponding posterior probability for evidence level defined by Pejaver et al. (2022). The intersection of those posterior lines represents the point at which a given REVEL or CADD score provides evidence towards either a benign (BP4) or pathogenic (PP3) classification. Fig3A shows the benign REVEL score thresholds, Fig3B shows the pathogenic REVEL score thresholds, Fig3C shows the benign CADD score thresholds, and Fig3D shows the pathogenic CADD score thresholds.

**Table 2.** CADD and REVEL evidence thresholds. The ClinVar 2019 score thresholds and score thresholds derived from a DVD likelihood with a DVD prior and tolerance-based prior for CADD or REVEL to provide evidence level for benign or pathogenic classification, as described in the text. *The Bin/Prior column uses either the DVD likelihood with tolerance-based bins or the DVD prior, or the score thresholds derived from the ClinVar 2019 dataset as defined in Table 1.

|  | Bin/Prior* | Benign |  |  |  | Pathogenic |  |  |  |
| --- | --- | --- | --- | --- | --- | --- | --- | --- | --- |
|  |  | Very Strong | Strong | Moderate | Supporting | Supporting | Moderate | Strong | Very Strong |
| CADD | Intolerant |  |  |  | 16.6 | 20.1 | 21.1 | 23.4 | 32.8 |
|  | Average Tolerance |  | 14.4 | 17.7 | 19.1 | 21.9 | 22.8 | 25.1 |  |
|  | Tolerant | 16.1 | 16.5 | 17.7 | 18.8 | 22.1 | 23.1 | 25.4 |  |
|  | X-linked/MT |  |  | 17.2 | 19.9 | 21.6 | 22.6 | 24.8 |  |
|  | DVD |  |  | 17.2 | 19.9 | 21.6 | 22.6 | 24.8 |  |
|  | ClinVar 2019 |  | 0.15 | 17.3 | 22.7 | 25.3 | 28.1 |  |  |
| REVEL | Intolerant |  |  | 0.050 | 0.138 | 0.267 | 0.338 | 0.495 | 0.826 |
|  | Average Tolerance | 0.087 | 0.110 | 0.178 | 0.237 | 0.375 | 0.451 | 0.620 | 0.947 |
|  | Tolerant | 0.096 | 0.122 | 0.194 | 0.255 | 0.396 | 0.475 | 0.646 | 0.954 |
|  | X-linked/MT | 0.073 | 0.097 | 0.164 | 0.223 | 0.359 | 0.433 | 0.600 | 0.938 |
|  | DVD | 0.073 | 0.097 | 0.164 | 0.223 | 0.359 | 0.433 | 0.600 | 0.938 |
|  | ClinVar 2019 | 0.003 | 0.016 | 0.183 | 0.290 | 0.644 | 0.773 | 0.932 |  |

Using strong levels of evidence, we calculated accuracy, sensitivity (ability to predict pathogenicity), and specificity (ability to predict benignity) for labeled DVD variants as summarized in Table 3. This analysis was repeated for ClinVar 2019 score thresholds and a DVD derived likelihood with a DVD prior or tolerance-based priors. Labeled benign and pathogenic variants that fall between the benign and pathogenic thresholds into what would be considered an uncertain range were counted as incorrectly labeled variants. Because the posterior for CADD did not reach a strong (posterior probability of 60%) evidence level for benign prediction in the analysis or in intolerant genes, a value of 0 was used in place of a lower threshold to calculate accuracy and specificity.

**Table 3.** Accuracy, Sensitivity and Specificity for CADD and REVEL with strong evidence of classification. Labeled benign and labeled pathogenic variants were used to evaluate ClinVar 2019 derived score thresholds and thresholds derived from the DVD likelihood and prior probability of 12.6% or tolerance-based prior unique to each bin defined in Table 1.

|  |  | Strong Evidence |  |  |
| --- | --- | --- | --- | --- |
|  | Prior | ClinVar 2019 | DVD | Tolerance Based |
| CADD | Accuracy | 0 | 0.095 | 0.549 |
|  | Sensitivity | 0 | 0.753 | 0.682 |
|  | Specificity | 0 | 0 | 0.526 |
| REVEL | Accuracy | 0.096 | 0.521 | 0.531 |
|  | Sensitivity | 0.345 | 0.792 | 0.723 |
|  | Specificity | 0.060 | 0.483 | 0.499 |

Overall, accuracy was highest when using the DVD likelihood and tolerance-based priors to define score thresholds with strong evidence from CADD at 54.9%, with similar accuracy from REVEL at 53.1%. REVEL (52.1% accurate) outperforms CADD (9.5% accurate) with a DVD prior due to the low specificity caused by the lack of strong evidence for benign prediction from CADD. The improved accuracy, sensitivity, and specificity from our method affirms the benefits of recalculating score thresholds using disease-specific datasets and binning genes by tolerance. While CADD accuracy is slightly higher than REVEL in the case of the tolerance-based prior, CADD cannot provide evidence for benignity in intolerant genes. Therefore, when using genetic tools alone, tolerance-based thresholds for REVEL have more consistent performance for strong evidence. The protein folding-informed analysis with CADD is included only in the supplemental data.

### Protein Folding-Informed Analysis

We predicted folding free energy differences (*ΔΔG*_*Fold*_) for all missense variants in the DVD to quantify their impact on protein stability. A *ΔΔG*_*Fold*_ greater than 1.0 kcal/mol leads to a five-fold increase in the ratio of unfolded to folded protein, indicating it is a reasonable threshold for variants that cause protein destabilization(29). There are 54,752 variants in the DVD that have a *ΔΔG*_*Fold*_ above 1.0 kcal/mol with 2,464 likely benign/benign, 2,884 likely pathogenic/pathogenic, and 49,404 VUSs.

We combined DVD derived likelihood ratios for *ΔΔG*_*Fold*_ and REVEL to determine the probability of a variant being pathogenic using both tolerance-based priors and a DVD prior. A posterior of 98% or greater is considered very strong evidence for pathogenicity(16). Using MATLAB contour and curve fitting functions, polynomial models were fit to define the posterior probability needed to reach each evidence level (see provided code). Details and the selection of prioritized variants are provided in the supplemental information.

**Figure 4.**
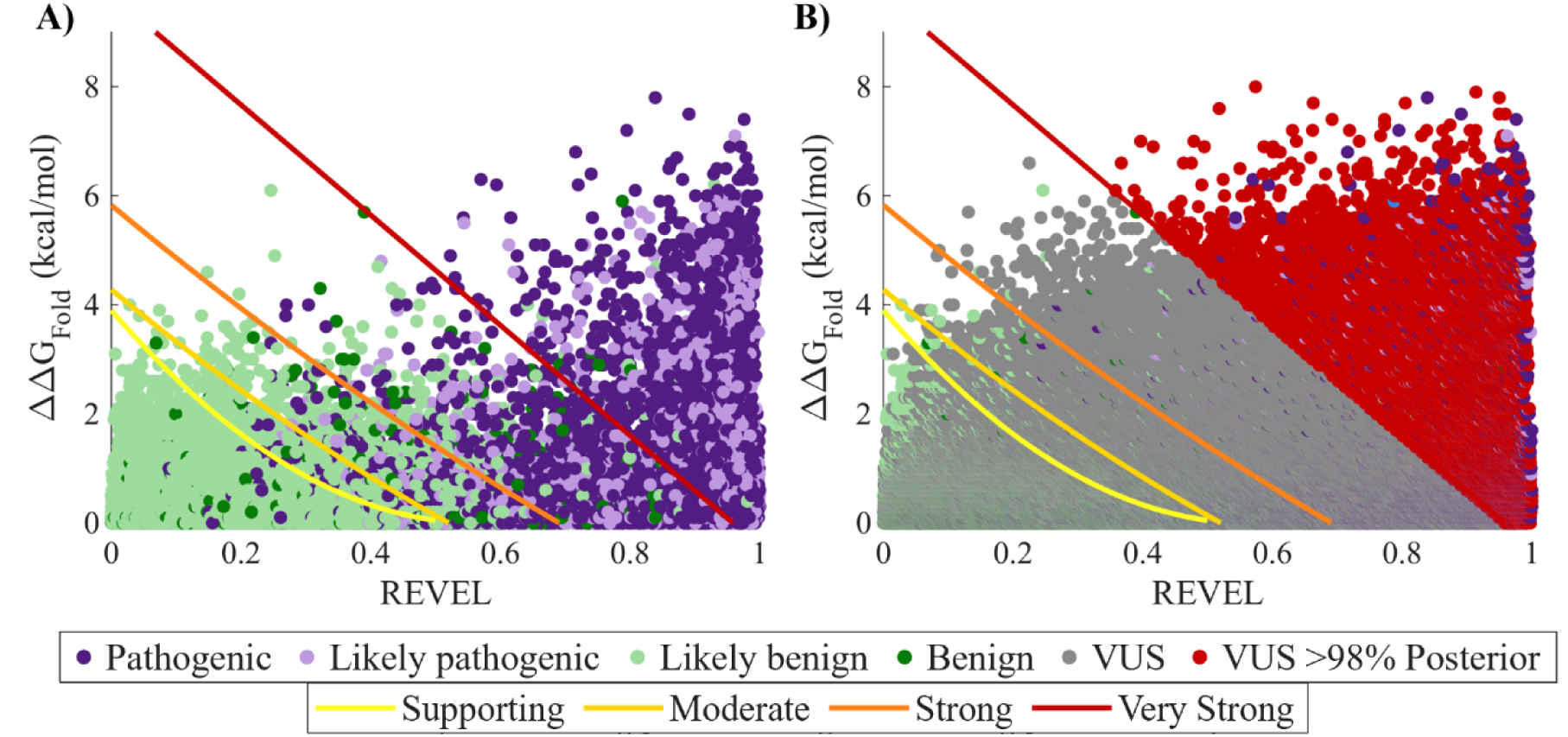
Scatterplots of ΔΔ*G*_*Fold*_ vs REVEL for missense variants with a DVD prior. Variants are colored by pathogenicity and the thresholds for evidence levels are shown. Fig4A shows only classified variants for REVEL x ΔΔ*G*_*Fold*_ with the four evidence thresholds. Fig4B shows REVEL x ΔΔ*G*_*Fold*_ with VUSs above 98% posterior probability colored red. A total of 18,706 of these VUSs exhibit a ΔΔ*G*_*Fold*_ greater than 1.0 kcal/mol.

The number of labeled variants and prioritized VUSs per tolerance-based bin for the ClinVar 2019 prior, the DVD prior, and the tolerance-based prior are summarized for REVEL x *ΔΔG*_*Fold*_ in Table 4. Results and figures for CADD x *ΔΔG*_*Fold*_ are outlined in the supplementary data.

**Table 4.** Variants with very strong evidence for pathogenicity. The classification of variants with very strong evidence from REVEL alone and REVEL x ΔΔ*G*_*Fold*_ using the ClinVar 2019 prior, the DVD prior, and the tolerance-based (TB) prior with the DVD derived likelihood. The + ΔΔ*G*_*Fold*_> 1.0 row shows the number of variants prioritized with a larger than 1.0 kcal/mol change in protein stability.

|  |  | Likely/Benign |  |  | Likely/Pathogenic |  |  | VUS |  |  |
| --- | --- | --- | --- | --- | --- | --- | --- | --- | --- | --- |
|  |  | ClinVar |  |  | ClinVar |  |  | ClinVar |  |  |
| Prior Odds |  | 2019 | DVD | TB | 2019 | DVD | TB | 2019 | DVD | TB |
| REVEL | Intolerant |  | 6 | 24 |  | 918 | 1,469 |  | 1,843 | 7,081 |
|  | Average Tolerance |  | 14 | 14 |  | 1,164 | 981 |  | 5,283 | 4,328 |
|  | Tolerant |  | 1 | 1 |  | 288 | 210 |  | 598 | 771 |
|  | X-linked/MT |  | 3 | 3 |  | 786 | 786 |  | 699 | 699 |
| REVEL x $\Delta\Delta G_{Fold}$ | Intolerant | 4 | 15 | 27 | 509 | 1,263 | 1,544 | 2,264 | 5,499 | 8,925 |
|  | Average Tolerance | 18 | 57 | 49 | 962 | 2,096 | 1,969 | 7,092 | 15,144 | 13,996 |
|  | Tolerant | 0 | 12 | 9 | 427 | 719 | 680 | 2,057 | 4,783 | 4,424 |
|  | X-linked/MT | 3 | 10 | 10 | 374 | 921 | 921 | 682 | 1,521 | 1,521 |
| **Total** | **REVEL** |  | **24** | **42** |  | **3,156** | **3,446** |  | **8,423** | **12,879** |
|  | **REVEL x $\Delta\Delta G_{Fold}$** | **25** | **94** | **95** | **2,272** | **4,999** | **5,114** | **12,905** | **26,947** | **28,866** |
|  | **+ $\Delta\Delta G_{Fold} > 1.0$** | **24** | **60** | **55** | **1,828** | **2,262** | **2,261** | **12,290** | **18,418** | **18,706** |

Application of the ClinVar 2019 and DVD prior has a false positive rate of 0.04% and 0.14% for REVEL x *ΔΔG*_*Fold*_, respectively. The tolerance-based DVD prior method also has a false positive rate of 0.14% (1 additional variant). Although the difference in false positive rate is minimal between all the priors, the tolerance-based prior correctly labels 2,842 or 115 additional pathogenic variants compared to the ClinVar 2019 and the DVD derived priors, respectively. It is difficult to calculate sensitivity (ability of a model to predict pathogenic variants) for the protein folding-informed thresholds since *ΔΔG*_*Fold*_ only tests one hypothesis for pathogenicity, and variants can be pathogenic due to other mechanisms (i.e. protein binding disruption). However, if we make the very liberal assumption that all labeled pathogenic variants not selected by our protein folding-informed thresholds are incorrectly labeled benign, the sensitivity for the ClinVar 2019, DVD, and tolerance-based protein folding-informed score thresholds are 23%, 50.7%, and 51.8%, respectively. Comparing REVEL x *ΔΔG*_*Fold*_ with REVEL very strong thresholds alone, the sensitivity is 32.0%, and the tolerance-based sensitivities are 34.9%. Our ability to predict pathogenic variants to a very strong evidence level is increased with protein folding-informed, tolerance-based priors with an increase of only 0.1% in benign variants being mislabeled when compared to the ClinVar 2019 prior and no increase compared to the DVD prior.

The tolerance-based protein folding-informed posterior assigns PP3 very strong evidence of pathogenicity to 28,886 VUS. The DVD protein folding-informed and ClinVar 2019 posteriors assign PP3 very strong evidence to 26,947 or 12,095 VUSs, respectively. The tolerance-based protein folding-informed posterior captures more VUSs than the DVD or ClinVar 2019 protein folding-informed methods with better sensitivity and a minimal increase in false positive rate. Figure 5 shows the score thresholds for REVEL x *ΔΔG*_*Fold*_ in the three bins. In Figure 5, Fig5A and Fig5B are the variants in the intolerant bin, Fig5C and Fig5D are those in the average tolerance range, and Fig5E and Fig5F are those in the tolerant bin colored by classification with VUS at or above a posterior of 98% colored red - see Figure 5).

**Figure 5.**
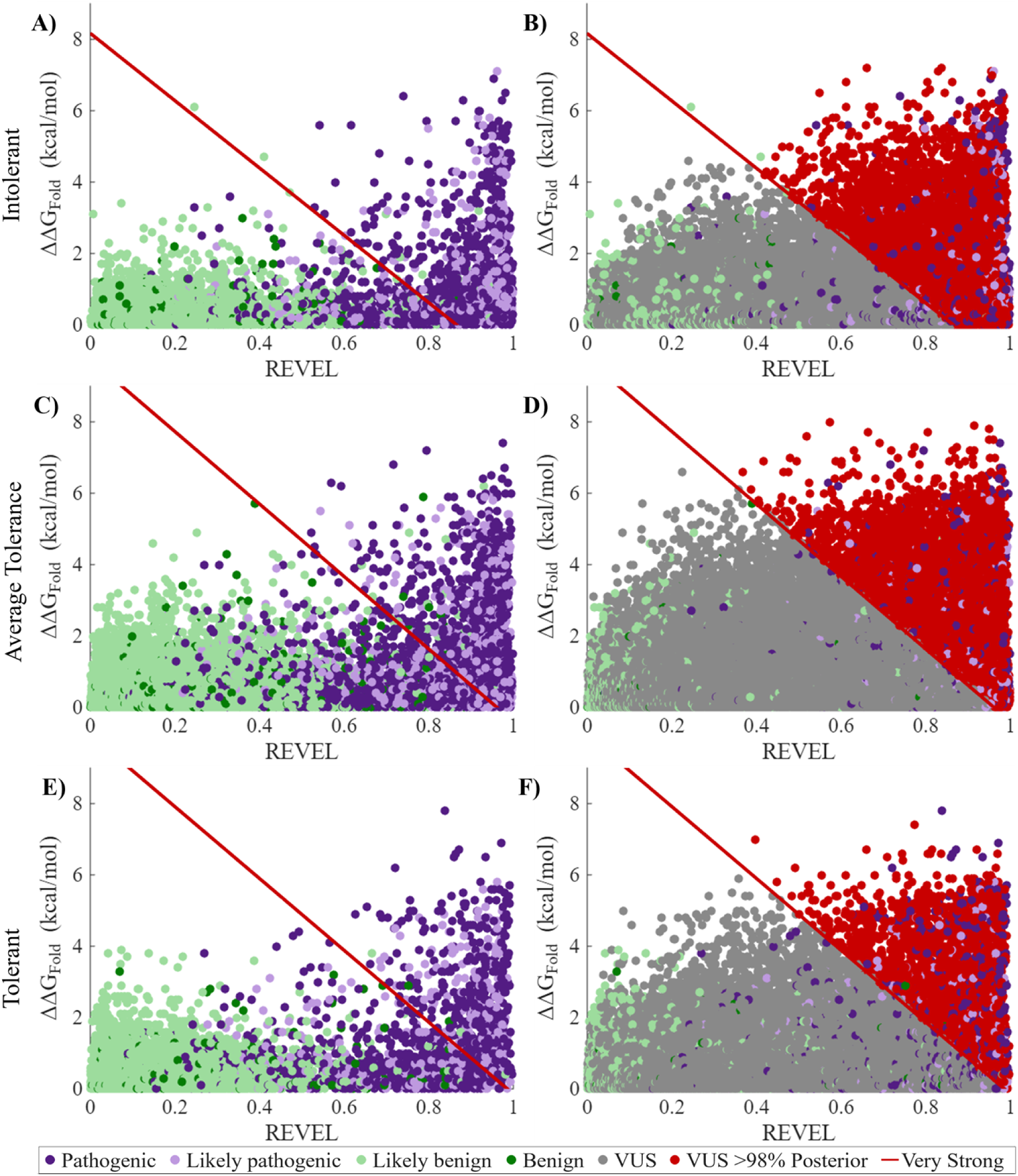
Scatterplots of REVEL x ΔΔ*G*_*Fold*_ for missense variants in tolerance-based bins. Variants are colored by pathogenicity and the thresholds for very strong in the intolerant, average tolerance, and tolerant bins are indicated by red lines. Fig5A shows classified variants for REVEL x ΔΔ*G*_*Fold*_ in the intolerant bin and Fig5B shows the REVEL x ΔΔ*G*_*Fold*_ score thresholds with prioritized variants colored red. Fig5C/D show REVEL x ΔΔ*G*_*Fold*_ show the same for the average tolerance bin and Fig5E/F for the tolerant bin.

Due to the significant increase in sensitivity, increase in VUS with PP3 very strong evidence of pathogenicity and low false positive rate, we recommend use of the REVEL x *ΔΔG*_*Fold*_ tolerance-based posterior for clinical use.

### Proband Variant Analysis

VUS with a 98% posterior probability of being pathogenic (based on the REVEL x *ΔΔG*_*Fold*_ tolerance-based posterior) were filtered against variants identified by OtoSCOPE^®^ v9 sequencing in probands with hearing loss to identify 12 variants. All 12 variants (outlined in the TableS3) received an upgraded genetic diagnosis to likely pathogenic/pathogenic with the inclusion of protein folding data, and 11 of the 12 variants have a clear structural mechanism to cause protein misfolding. Two cases detailed below illustrate the impact of these variants on protein structure and misfolding (Fig. 6).

**Figure 6.**
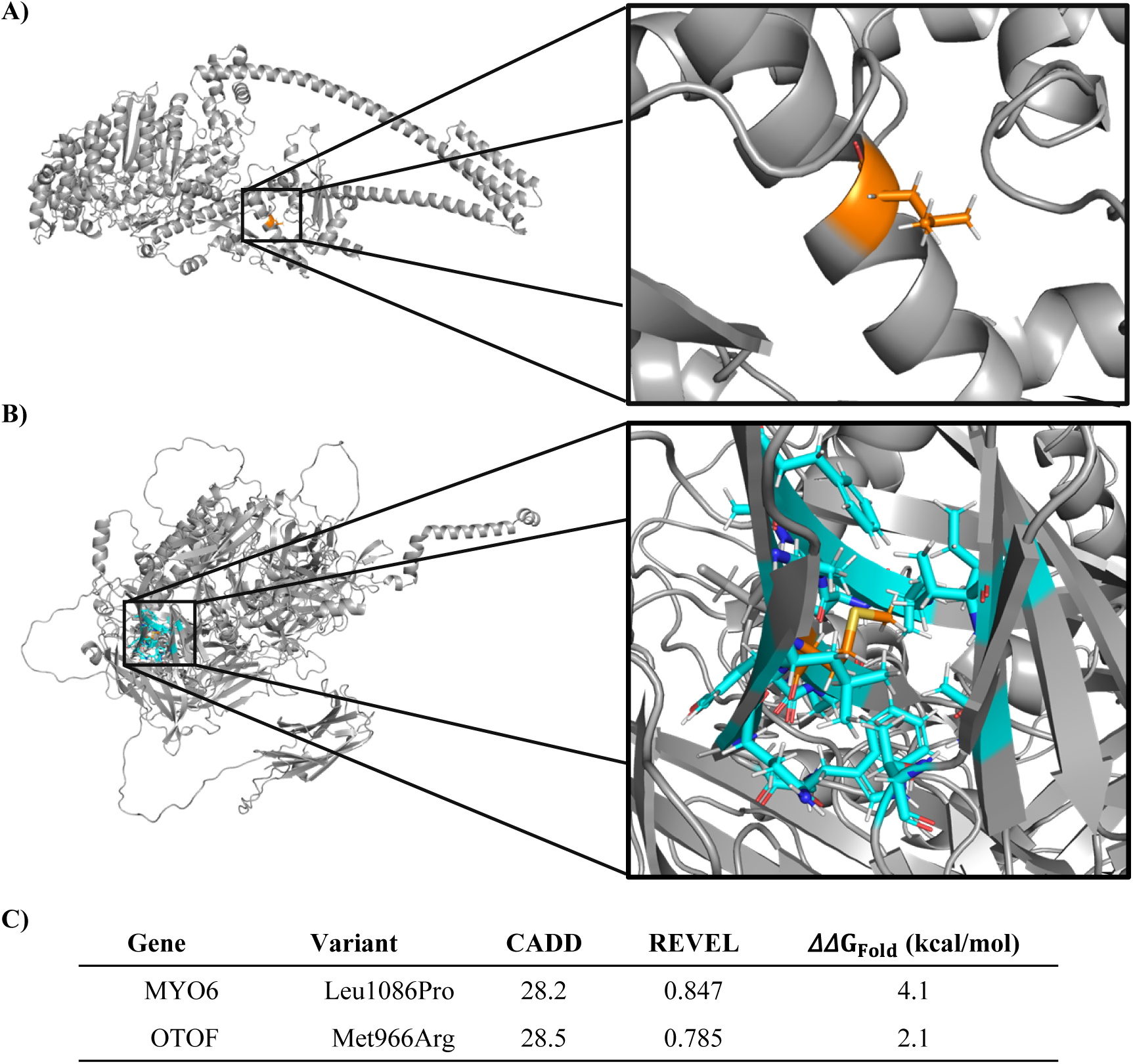
Highlighted patient variants in their corresponding full-length protein structures. Fig6A shows the full-length *MYO6* protein structure with the variant p.(Leu1086Pro) and the residue Leu1086 colored orange. Fig6C shows the full-length *OTOF* protein structure with the variant p.(Met966Arg) where the residue Met966 is colored orange and the residues surrounding hydrophobic residues are colored cyan. Fig6D gives the CADD, REVEL, and *ΔΔG*_*Fold*_ score for each variant.

### *MYO6* p.(Leu1086Pro)

Variants in *MYO6* are associated with autosomal dominant non-syndromic hearing loss(30, 31). The encoded protein, myosin VI (MyoVI), is an unconventional actin-based motor important for endocytosis in cochlear hair cells(32). A sib pair with congenital mild-to-moderate bilateral sensorineural hearing loss were heterozygous for the *MYO6* variant, p.(Leu1086Pro). This variant has a REVEL score of 0.847 and causes a destabilizing *ΔΔG*_*Fold*_ of 4.1 kcal/mol. Proline is unique among amino acids due to the covalent bond its side chain forms with the protein backbone, which makes it relatively rigid and prevents formation of secondary structure due to the inability of the backbone nitrogen atom to donate a hydrogen bond. As residue Leu1086 is in the middle of an alpha helix (Fig6A), the helix is broken by the introduction of a proline, significantly impacting the structure and stability of the protein

### *OTOF* p.(Met966Arg)

*OTOF* encodes for otoferlin, a calcium binding transmembrane protein in synaptic vesicles important for signal transmission in inner hair cells(33, 34). Mutations in this gene lead to autosomal recessive severe-to-profound bilateral hearing loss. Accurate classification of variant effect is of prime importance as there are now several gene therapy options for persons with *OTOF*-related hearing loss(35–38). OtoSCOPE^®^ identified the heterozygous *OTOF* variant, p.(Met966Arg) in a proband with early childhood onset profound hearing loss and a family history of hearing loss in a sibling. Importantly, this proband was heterozygous for the known pathogenic *OTOF* variant, p.(Ile1573Thr). *OTOF* has a missense o/e score of 0.94, falling within average tolerance. The variant p.(Met966Arg) has a REVEL score of 0.785 and a *ΔΔG*_*Fold*_ of 2.1 kcal/mol indicating a destabilizing effect on the protein folding. Methionine is a hydrophobic residue, and the residue Met966 (orange in Fig6C) is surrounded by other hydrophobic residues (cyan in Fig6C). The hydrophobic effect describes the thermodynamically unfavorable interaction of hydrophobic protein residues with water, which drives protein folding by the favorable burial of hydrophobic residues into a hydrophobic protein core as seen in Fig6C(39). Arginine is positively charged and hydrophilic, which disrupts the stabilizing hydrophobic core consistent with the high

*ΔΔG*_*Fold*_.

## Discussion

Although genome sequencing offers the promise of personalized precision medicine, variant classification remains a challenge. In the domain of hearing loss, for example, there are 381,924 missense variants in the 224 genes represented on the OtoSCOPE v9 comprehensive genetic testing panel – 303,577 of these variants (79.5%) are VUSs. Generating experimental evidence to phenotype these variants is implausible, highlighting the importance of optimizing variant analysis using *in silico* tools. Here, we construct a Bayesian framework for evaluating VUSs to define deafness-specific thresholds augmented with protein folding modeling to identify VUSs in the DVD that have a 98% posterior probability of being pathogenic(16).

We refined the structure of 258 unique protein isoforms for 224 genes in the DVD predicted by AlphaFold3 with the AMOEBA force field to ensure local structural minimization and global side chain optimization. Using these optimized structures (OtoProteinV3), we predicted folding free energy differences (*ΔΔG*_*Fold*_) for all missense variants in the DVD. We then evaluated REVEL, CADD, and *ΔΔG*_*Fold*_ by applying a Bayesian framework with a tolerance-based prior specific for hearing loss genes(16). We found that a prior probability of pathogenicity of 12.6% (specific for hearing loss) increased accuracy, sensitivity, and specificity for REVEL scores when compared to ClinVar 2019 derived thresholds (ClinVar 2019 CADD thresholds could not provide strong or very strong evidence of pathogenicity; *ΔΔG*_*Fold*_ was not predictive on its own with a 4.3% accuracy). When using tolerance-based thresholds, accuracy and specificity were improved for both CADD and REVEL, although sensitivity decreased when compared to using the DVD prior.

The choice of prior probability has a profound effect on deleteriousness predictions: a low prior probability of 4.1% (prior odds 0.046) as used by Pejaver *et al.* for predictions on the ClinVar 2019 dataset under calls pathogenic variants in the context of the DVD. The ClinVar 2019 prior is also significantly lower than previous estimates including those defined by other work on deafness phenotypes(27). Use of a DVD prior of 12.6% (prior odds 0.144) is more consistent with labeled DVD variants based on improved accuracy, sensitivity and specificity. Our tolerance-based bins suggest that highly constrained genes such as *ACTG1* require an even larger prior (25.7%) to explain the observed data. We next calculated likelihoods for CADD and REVEL with *ΔΔG*_*Fold*_ to combine biophysical evidence with genetic tools. Using a tolerance-based prior and REVEL x *ΔΔG*_*Fold*_ likelihood resulted in score thresholds with a false positive rate of 0.14%, a sensitivity of 51.8% and identify 28,886 of 303,577 (9.5%) variants with a 98% posterior probability of being pathogenic. A total of 18,706 of the VUS identified have a protein folding free energy difference greater than 1 kcal/mol, which indicates around two thirds of the VUS captured by the protein folding-informed thresholds are implicated due to their downstream effects on protein structure and stability. This is consistent with previous studies that conclude approximately ∼60% to ∼83% of disease-causing missense variants result in loss of protein stability(19–21). When looking at already labeled pathogenic/likely pathogenic variants, 2,884 variants have a protein folding free energy difference above 1.0 kcal/mol. Practically, adopting 10% as a conservative minimum as recommended by the ClinGen Structural Variant Interpretation working group, labels approximately 40,000 variants as LP/P (currently labeled LP/P values together with VUS above 98% posterior probability), i.e., the roughly 28,000 upgraded VUS are consistent with expectations(27). With a false positive rate of 0.1–0.2%, we would expect fewer than 100 erroneous promotions—arguably an acceptable tradeoff for greater sensitivity.

The DVD is resource that is regularly updated, experiencing a nearly 4-fold increase in missense variants from 2023 to 2026. Calculating the prior probability rate of pathogenicity for the previous instance of the DVD (v8) yields nearly 30%, which is much higher than the value for the current version of 12.6%. As more variants are labeled by the field, the prior rate will converge toward the “correct” value, including additional support for tolerance-based or gene specific models. The classifications in the DVD will also continue to improve as the ACMG/AMP criteria evolve and the internal evaluation of variants advances.

The clinical impact of this data is significant. In investigating variants in probands sequenced with OtoSCOPE, we used the protein folding-informed thresholds to upgrade 12 VUS to likely pathogenic/pathogenic. As an example, we highlight two variants identified in persons with hearing loss – *MYO6* p.(Leu1086Pro), and *OTOF* p.(Met966Arg) – and show how close structural analysis of protein folding contributes to our understanding of variants. *MYO6* p.(Leu1086Pro) causes a disruption to folding of the alpha helix due to proline rigidity; *OTOF* p.(Met966Arg) disrupts the hydrophobic pocket secondary to arginine positive charge, thereby affecting the overall fold.

There are important limitations to this work: 1) the possibility of overestimating the performance due to circularity in variant classification data, 2) the limited ability of the methods to predict benign variants, and 3) limitations inherent to each computational tool (CADD, REVEL, AlphaFold3 and DDGun3D). First, some DVD variants were included in the original training sets used for CADD and REVEL, which were not removed for the current assessments and can artificially inflate performance. Additionally, the variant classification of likely benign is assigned to some variants in the DVD using a custom bioinformatics pipeline. This pipeline utilizes computational tools to classify variants and introduces artificial inflation of the benign estimates in the DVD. We also introduce some overestimation when combining the likelihoods of CADD or REVEL with *ΔΔG*_*Fold*_. Bayesian statistics stipulate that to combine likelihoods by multiplication the variables must be independent. CADD or REVEL and *ΔΔG*_*Fold*_ are not entirely independent (see Supplemental Data) with correlation coefficients 0.08 and 0.16, respectively. However, the relatively small correlation coefficients together with the low false positive rates support the conclusion that the correlation is minor and does not hinder our protein folding-informed models.

Second, CADD does not reach strong evidence for pathogenic or benign variant classification. When coupled with the overall lower specificity observed for all methods of predicting labeled benign variants, these recalibrated thresholds lack sufficient power to reliably detect benign variants.

Third, the training set for AlphaFold3 is from experimental structures available in the Protein Databank. For approximately 60% of the proteins in this work, no experimental or homology-related structural information was available for AlphaFold2 or 3(18, 40, 41). There remain inaccuracies for a subset of the predicted structural models that could impact the accuracy of predicted *ΔΔG*_*Fold*_ values. We expect more accurate protein structure predictors (Boltz-2(42, 43), OpenFold(44, 45)) that will overcome the limitations of AlphaFold3. While DDGun3D was used in this study due to its combination of accuracy and efficiency, alternative methods for the prediction of protein folding free energy differences are emerging(46). Ideally, we would calculate *ΔΔG*_*Fold*_ with rigorous molecular dynamics simulations, however, this option is currently not feasible at scale for more than 380,000 missense variants. We expect future machine learning and/or physics-based methods to emerge that will approach the accuracy of *in vitro* folding experiments.

As more accurate prediction methods for both protein structure and *ΔΔG*_*Fold*_ become available, we will continue to investigate all variants in the DVD(47). Molecular simulation methods for quantifying *ΔΔG*_*Fold*_ will become faster and allow for thorough assessment of biophysical phenotypes(48, 49). Structure prediction methods will continue to evolve and allow for the creation of protein-protein complexes, protein-ligand interactions, and the inclusion of RNA/DNA. Each of these structures has different quantifiable evidence (*e.g.*, like ΔΔG_Fold_ for misfolding) that can be used to evaluate the mechanism of disease for missense variants(50, 51). It is realistic to expect future work to include binding free energy differences for protein-protein, protein-ligand, and protein-DNA interactions. In the future, we believe *in silico* tools will be able to provide adequate evidence for variant classification.

It is important to note that ACMG/AMP guidelines stipulate that no *in silico* tool provides adequate evidence to classify a single variant as benign or pathogenic. Therefore, computational evidence must be integrated with other lines of evidence, such as functional or population data, before a final classification can be made(6). For that reason, the DVD will include the posterior probability for all missense variants, which contributes PP3 evidence of various strengths (*i.e.*, very strong when the posterior is greater than 98%) for clinical interpretation.

This work also indicates there is additional evidence in mutation tolerance scores from gnomAD that can inform our investigation of gene-specific variants in the DVD. The nature of this impact is indicated by the increased prior odds for intolerant genes and the reduced prior odds for tolerant genes. ACMG PP2 criteria state that missense variants have additional evidence for pathogenicity if they occur within genes where few missense variants are benign and missense variation is a common mechanism of disease. Our intolerant prior captures and quantifies this information to improve our prediction of pathogenic variants. While this could lead to over-counting evidence if both PP2 and our evidence is used when evaluating variants, our intolerant priors also allow for additional information in genes where there are too few labeled variants to apply PP2. Tolerance information will aid clinicians in evaluating the impact of novel missense variants as the prior probability of pathogenicity varies greatly (e.g., from more than 25% in intolerant genes down to below 10%) (6).

In conclusion, deafness-specific Bayesian models improve the accuracy of REVEL and CADD for missense variant classification in hearing loss by increasing sensitivity for pathogenicity prediction. While limitations remain for the classification of benign variants, these results underscore the importance of disease-specific thresholds. Combining genetic-based predictors of variant effects that rely heavily on evolutionary conservation with biophysical evidence allows us to better prioritize VUS as pathogenic. Use of protein folding free energy differences directly tests a mechanistic hypothesis for the underlying biophysical impact of each missense variant, unlike CADD or REVEL that fail to provide a phenotypic rationale for their score. We define a Bayesian framework using a tolerance-based prior and likelihood that combines REVEL with *ΔΔG*_*Fold*_ to assign over 28,000 VUS with PP3 very strong evidence (i.e., 98% posterior probability in favor of pathogenicity). Finally, while the methods described here are applied to hearing loss, the pipeline for variant evaluation could be applied in other disease contexts.

## Material and Methods

### Dataset

Genes and variants associated with hearing loss or deafness have been compiled in the publicly available Deafness Variation Database (DVD) (https://deafnessvariationdatabase.org) (7). Variants are collected from dbSNP(8), gnomAD(9), the 1000 Genomes Project(10), ClinVar(11) and HGMD(12). In total, the DVD contains more than 10.5 million variants. Here, we focus on 381,924 missense variants across 216 DVD genes that cause a downstream single amino acid change (the remaining 8 genes do not have missense variants: *MT-RNR1, MT-TH, MT-TI, MT-TK, MT-TL1, MT-TS1, MT-TS2, MIR96*).

### AlphaFold3

AlphaFold3(18) is an updated version of the deep learning structure prediction model AlphaFold2(40) that contains a diffusion-based architecture to predict protein monomer structures. Previous work using the AlphaFold2 network predicted protein structures for all DVD genes and their corresponding isoforms referred to as OtoProteinV2(23). AlphaFold3, however, offers improved accuracy over AlphaFold2, motivating updated protein models(18). For example, the DVD includes several proteins with sequences larger than 2,400 amino acids; these proteins were predicted in sections and pieced together manually in OtoProteinV2. The AlphaFold3 webserver offers the capability to model proteins up to 5,000 amino acids, reducing the cost of predicting the larger DVD protein structures and clashes introduced from stitching together isolated protein structural domains. Three genes (ADGRV1, KMT2D, USH2A) in the DVD encode extremely large proteins (above the 5,000 amino acid limit of AlphaFold3). For these proteins, we transferred the OtoProteinV2 model into this dataset but improved on the original optimization. All DVD genes and their relevant isoforms have full-length protein models.

### Optimization of Monomer Structures

AlphaFold3 performs a local minimization in the Amber fixed-charge force field as part of the prediction(52, 53). Even with Amber optimization, the structures have atomic van der Waals clashes and high-energy backbone torsions that can be improved using local backbone optimization together with global side chain optimization(54, 55) based on the **A**tomic **M**ultipole **O**ptimized **E**nergetics for **B**iomolecular **A**pplications (AMOEBA) polarizable force field(56, 57). Initial local minimization was performed with the L-BFGS optimization algorithm to a convergence criterion of 0.8 kcal/mol. Protein models then underwent global side chain optimization before a final local minimization was performed to a convergence criteria of 0.1 kcal/mol.

We evaluated the structures before and after AMOEBA optimization in Force Field X(58) with MolProbity, a tool that scores protein structures based on a database of well understood conformational features (e.g., equilibrium bond lengths, protein backbone geometry, and side chain rotamers). A MolProbity score of 1.0 implies the expected quality of a structural model from X-ray diffraction at a resolution of 1.0 Å(59, 60), whereas higher scores indicate lower quality models consistent with lower quality diffraction data. The final optimized protein structures are available on GitHub (https://github.com/SchniedersLab/OtoProtein3) and through the DVD website.

### Predicting ΔΔG_Fold_

We used a regression based *ΔΔG*_*Fold*_ predictor called DDGun3D(46) to evaluate missense variants in protein structures. DDGun3D is a high throughput predictor for *ΔΔG*_*Fold*_ that uses a linear regression of biochemical and structural features to determine the thermodynamic effects of missense variants(46). DDGun3D has a reported root-mean-square-error (RMSE) of ∼1.5 kcal/mol. The advantages of DDGun3D over other prediction methods are its ability to be applied to a dataset of our scale with minimal compute cost and the anti-symmetry of the method. Perfect anti-symmetry constrains the free energy change of mutating from residue A→B to be equal and opposite in sign to the free energy change of mutating from B→A. DDGun3D was used in a previous study on the DVD(23) and applied here on the updated 381,924 missense variants in the current DVD instance.

### Bayesian Analysis

To determine score thresholds consistent with ACMG/AMP evidence levels, we used an established Bayesian framework that focused on evaluating separate tools for their ability to predict variant pathogenicity(16). ClinVar 2019 derived score thresholds for REVEL and CADD are reported with varying levels of evidence (supporting, moderate, strong, and very strong) to indicate that a variant is pathogenic or benign. To determine disease-specific score thresholds, we applied this framework to a dataset of well-curated variants from the DVD.

We next calculated a pathogenic and benign likelihood ratio for each genetic tool (CADD, REVEL, and *ΔΔG*_*Fold*_). *ΔΔG*_*Fold*_ has limited ability to predict classifications on its own, however, we include *ΔΔG*_*Fold*_ applied in isolation in the supplementary data.

First, the prior odds of pathogenicity for the DVD were calculated.

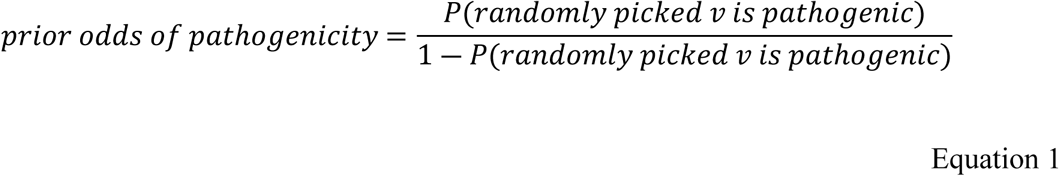

where *v* is a variant. Pejaver reported the prior probability of pathogenicity as 4.1% (prior odds of 0.046) based on the ClinVar 2019 dataset. This is below the ClinGen Structural Variant Interpretation working group recommendation of a prior probability of pathogenicity of ∼10%(26). The prior probability of pathogenicity for the DVD based on currently classified variants (32,895 benign, 65,585 likely benign, 3,205 likely pathogenic, and 6,662 pathogenic) is 12.6% (prior odds of 0.144). The higher prior is consistent with selection of OtoSCOPE^®^ genes with known disease associations and their relative (in)tolerance to missense variants (as discussed further in the next section on gene tolerance). We then calculate the likelihood ratio for a given score using Equation 2.

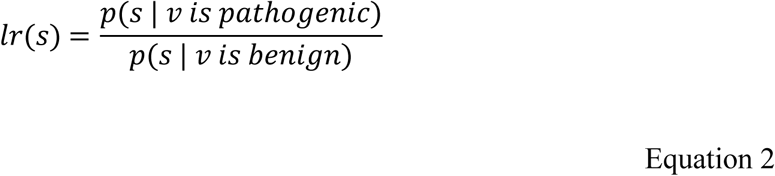

where *s* is the score (CADD, REVEL, or *ΔΔG*_*Fold*_). The likelihood ratio for each metric is calculated for 10 discrete bins that span the scoring range of each metric and ensures there are enough classified variants within each bin to calculate a reliable likelihood. The likelihood ratios for each tool are in TableS1. The likelihood for CADD or REVEL was combined with the likelihood for *ΔΔG*_*Fold*_ to define protein folding-informed (PFI) likelihoods where *lr*_*PFI*_ = *lr*_*CADD*/*REVEL*_ ∗ *lr*_ΔΔ*GFold*_. Equation 3 is applied to *lr*_*CADD*_, *lr*_*REVEL*_, and *lr*_*PFI*_.

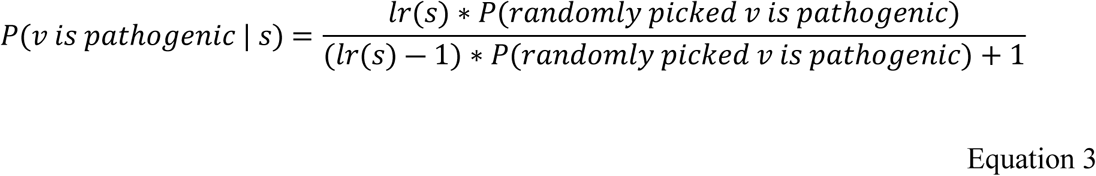

where *P*(*v is pathogenic* | *s*) is the posterior probability a variant is pathogenic given some score. Posterior values of 98% correspond to PP3 very strong evidence for pathogenicity(16).

We determine the CADD or REVEL score threshold for the posterior to reach supporting (10%), moderate (20%), strong (60%), and very strong (98%) evidence(16). It is important to note variants that were included in the training set for CADD and REVEL were removed from the ClinVar 2019 dataset to avoid biasing the thresholds for evidence types. For this study, we included all variants to fully understand the impact on the DVD. We used the calculated disease-specific score thresholds and compared them to those established in with the ClinVar 2019 dataset for their ability to predict labeled variants in the DVD.

To calculate the protein folding-informed score thresholds for REVEL in combination with *ΔΔG*_*Fold*_, we calculated the posterior for all combinations of REVEL and *ΔΔG*_*Fold*_ likelihoods (*i.e.*, for 10 x 10 = 100 bins). Variants from REVEL x *ΔΔG*_*Fold*_ bins above the 98% posterior threshold (very strong evidence) were added to a prioritized list for the protein folding-informed score threshold. We performed the same analysis with CADD and *ΔΔG*_*Fold*_ with results outlined in the supplementary data.

### Checking for independence between tools

We performed a regression analysis on DDGun3D values vs CADD/REVEL to determine if the variables are independent, which is necessary to justify creation of a combined likelihood as the product of each tool’s individual likelihood. The R-squared coefficient between DDGun3D and CADD/REVEL was just 0.08 and 0.16, respectively. For two scores to be completely independent, their correlation would be 0. Although the correlation coefficient is slightly above 0, the low false positive rate of the combined likelihoods supports the approximation of treating CADD/REVEL and DDGun3D as independent in defining their joint likelihood ratio.

### Prior Probability from Genes Grouped by Tolerance

For each gene in the DVD, we collected a missense tolerance score from gnomAD as a means to create groups (bins) of genes whose missense variants were hypothesized to have similar prior probabilities of being pathogenic(61). The tolerance score, missense o/e, is a ratio of the observed-to-expected number of variants in a particular gene. The observed variant count is the number of unique single nucleotide variants in the transcript with minor allele frequency of less than 0.1% and a median depth in exome samples greater than 30. The expected variant count uses a model that corrects for local sequence context as described in Karczewski *et al*(22). The final ratio provides a continuous measure of gene tolerance to mutation where higher scores are more tolerant and lower scores are less tolerant (or more intolerant). We hypothesized that genes with a lower o/e (more intolerant) score would have a higher prior probability of pathogenicity(62).

We used missense o/e scores to bin genes in the DVD based on their tolerance and calculated priors based on the genes/variants in each bin. Genes in the DVD had missense o/e values ranging from 0.1 (intolerant) to 1.45 (tolerant) and were divided into three separate bins. The missense o/e and genes/variants in each bin are shown in Table 1 along with their prior probability of pathogenicity and benignity. The bins are not evenly distributed by o/e score; the first and last bins are significantly larger than the middle bin to capture the less common extremes of the o/e score. The *average* tolerance (middle) bin is defined by using the mean missense o/e as an approximate center and includes the central distribution of the DVD. The two outer bins (tolerant and intolerant) contain the extreme ends of the distribution. There are 16 genes that are X-linked or mitochondrial (X-linked/MT) and do not have an o/e score per gnomAD. The X-linked/MT genes were assigned the overall DVD prior. We compared the accuracy, sensitivity, and specificity when using tolerance-based priors to using a single DVD prior with the DVD derived likelihoods to establish score thresholds and the score thresholds derived from the ClinVar 2019 dataset to select a genetic tool and protein folding-informed Bayesian framework for evaluating variants.

### Proband Variant Analysis

As a proof of concept, to understand the translational impact of our protein folding-informed Bayesian framework, we identified probands sequenced on OtoSCOPE^®^ v9 panel without a definitive genetic diagnosis but with at least one VUS within the group of more than 28,000 VUSs prioritized by REVEL x *ΔΔG*_*Fold*_ to be pathogenic. Matched variants received additional scrutiny (a rigorous structural and genetic analysis) to determine their likely effect on protein structure/function.

## Acknowledgements

Author RAG was supported by NIH T32 Grant No. 2T32GM008365-26A 1, the Center for Biocatalysis and Bioprocessing, NIH R01DC012049, and the Graduate College Iowa Recruitment Fellowship. Author MJS was supported by NIH R01DC012049 to study the roles of protein misfolding on hearing loss and NSF CHE-2504153 for the development of the protein structure optimization and thermodynamics methods leveraged in this work. Author RJHS was supported by NIH R01DC002842, R01DC012049, and R01DC017955. Author TAB and GMC were supported by NIH R01DC012049.

## Author Contributions

RAG, GMC, MJS, and TAB conceived the idea. MJS and TAB supervised the study. RAG and GMC designed the model. RAG and CEO performed structural analysis. RAG and GMC performed Bayesian analysis. AMO, KLM, and RAG performed proband analysis. RJHS, AMO, KLM, BH, and DLK provided project support. HA and TLC reviewed and revised the manuscript. All authors have revised and approved the manuscript.

## Data Availability

The dataset used to complete this study is freely available on the DVD website (https://deafnessvariationdatabase.org). AlphaFold3 is available as a webserver and Force Field X is available at https://ffx.biochem.uiowa.edu. Computational tools used in this study (CADD, REVEL, DDGun3D) are freely available. The protein structural models are available on the DVD website and on GitHub at https://github.com/SchniedersLab/OtoProtein3. The biophysics data, o/e scores, and posterior probabilities for all variants will be integrated into the DVD website and are included as a supplementary data file. The selected VUS are included as a supplementary data file. The MATLAB scripts used to analyze data are included as supplementary files.

